# Multiple overlapping dynamic patterns of the visual sensory network in schizophrenia

**DOI:** 10.1101/2020.12.21.423535

**Authors:** Mohammad S. E Sendi, Godfrey D. Pearlson, Daniel H. Mathalon, Judith M. Ford, Adrian Preda, Theo G. M. van Erp, Vince D. Calhoun

## Abstract

Although visual processing impairments have been explored in schizophrenia (SZ), their underlying neurobiology of the visual processing impairments has not been widely studied. Also, while some research has hinted at differences in information transfer and flow in SZ, there are few investigations of the dynamics of functional connectivity within visual networks. In this study, we analyzed resting-state fMRI data of the visual sensory network (VSN) in 160 healthy control (HC) subjects and 151 SZ subjects. We estimated 9 independent components within the VSN. Then, we calculated the dynamic functional network connectivity (dFNC) using the Pearson correlation. Next, using k-means clustering, we partitioned the dFNCs into five distinct states, and then we calculated the portion of time each subject spent in each state, that we termed the occupancy rate (OCR). Using OCR, we compared HC with SZ subjects and investigated the link between OCR and visual learning in SZ subjects. Besides, we compared the VSN functional connectivity of SZ and HC subjects in each state. We found that this network is indeed highly dynamic. Each state represents a unique connectivity pattern of fluctuations in VSN FNC, and all states showed significant disruption in SZ. Overall, HC showed stronger connectivity within the VSN in states. SZ subjects spent more time in a state in which the connectivity between the middle temporal gyrus and other regions of VNS is highly negative. Besides, OCR in a state with strong positive connectivity between middle temporal gyrus and other regions correlated significantly with visual learning scores in SZ.

## 1. Introduction

Schizophrenia is a common, chronic, developmental brain disorder thought to involve aberrant connectivity (Weinberger, 1988). In recent years, functional connectivity data derived from resting-state functional magnetic resonance imaging (fMRI) time series has proven highly informative regarding underlying brain connectivity patterns in schizophrenia (Adhikari et al., 2019; Hummer et al., 2020). The disease has been characterized as one of “brain dysconnectivity” (Friston, 1998). More recently work has focused on the dynamics of functional network connectivity or dFNC (Damaraju et al., 2014; Dong et al., 2019; Garrity et al., 2007; Miller et al., 2016; Sendi et al., 2020; Sun et al., 2019). Visual processing impairment is a known problem in SZ, and the visual sensory network (VSN) likely plays a key role in the disorder. However, there has not to date been a focused study on the resting fMRI dynamics among visual related regions in schizophrenia.

Sensory processing dysfunction, including visual sensory, was reported in schizophrenia (Javitt, 2009), as was an impairment of perceptual closure and its positive correlation with symptom severity (Van de Ven et al., 2017). The latter study also reported a reduction in visual cortex intrinsic connectivity. A reduction in the P1 component of the visual event-related potential highlighted an early visual processing deficit in schizophrenia subjects and their relatives (Yeap et al., 2008, 2006), suggesting that this may be an endophenotype.

All of the above studies show the potential role of the visual processing regions as an underlying mechanism in schizophrenia. Also, dFNC is informative in studying the underlying mechanisms of brain function in many diseases (Damaraju et al., 2014; Dong et al., 2019; Fiorenzato et al., 2019; Garrity et al., 2007; Miller et al., 2016; Sun et al., 2019; Zhi et al., 2018). Thus, we hypothesized that studying the dynamics of the visual network may be informative about in what manner and the degree to which the visual system is disrupted in schizophrenia. We expect that the state-dependent connectivity disruption within a shorter timescale would reveal more information about the dynamics of VSN in schizophrenia, and potentially the investigation of the link between visual learning and VSN dFNC provides additional insight into visual learning impairment in SZ subjects.

In this study, we leveraged the sliding window approach followed by *k*-means clustering to identify a set of connectivity states to investigate dFNC within the visual network (Allen et al., 2014). To further investigate and model these temporal changes, we estimated the occupancy rate (OCR) of each state from dFNC. Next, via statistical analysis on the estimated OCR features, we explored the difference between schizophrenia and healthy subjects. In addition, we compared cell-wise differences between SZ and HC within each state. Finally, we investigated the link between OCR features and visual learning in SZ subjects.

## 2. Methods

### 2.1. Participants and dataset

We used rs-fMRI and clinical data from the Functional Imaging Biomedical Informatics Research Network (FBIRN) projects (Van Erp et al., 2015). The raw imaging data were collected from seven sites including the University of California, Irvine; the University of California, Los Angeles; the University of California, San Francisco; Duke University/the University of North Carolina at Chapel Hill; the University of New Mexico; the University of Iowa; and the University of Minnesota. The written informed consent, approved by institutional review boards of each study site, was obtained from all subjects. All patients were on stable doses of antipsychotic medication, either typical, atypical, or a combination for at least 2 months at the time of the study (Keator et al., 2016). Stable medication means that there was not a significant change in the medication does of each SZ subject for two months before the scanning time. All SZs were clinically stable at the time of scanning. A diagnosis of schizophrenia was confirmed using the SCID-IV interview (First et al., 2002a) and an absence of schizophrenia diagnosis in HC was confirmed with the SCID-I/NP interview (First et al., 2002b). In addition, HC subjects with a current and past history of major neurological or psychiatric disorder and with a first-degree relative with an Axis-I psychotic disorder diagnosis were also excluded assessed by SCID.

The imaging data were collected with six 3T Siemens and one 3T General Electric scanner. All scanning sites used the same protocol for collecting the rs-fMRI. T2*-weighted functional images were collected using AC-PC aligned echo-planar imaging sequence with TE=30ms, TR=2s, flip angle = 77°, slice gap=1 mm, voxel size= 3.4 × 3.4 × 4 mm^3^, and 162 frames, and 5:24 min. All participants were instructed to close their eyes during the rs-fMRI data collection.

### 2.2. Visual learning

All participants had normal eyesight and were fluent in English. The visual figure learning test (VFLT) from the computerized multiphasic interactive neurocognitive diagnostics system (CMINDS) was used to test the visual learning in the participants (O’Halloran et al., 2008; Van Erp et al., 2015) and was performed outside of the scanner. Six geometric figures in a 3×2 matrix were presented to participants for 10 seconds. Then, subjects are asked to draw as many of the presented figures as they can recall on a blank screen in the same location as they were presented originally. Around 25 minutes after completing the immediate recalling test, the subjects are asked to reproduce as many of the 6 original figures as they can recall. Next, 12 figures, including 6 original and 6 new, are presented to the subjects, and they were asked to answer “No” if the figure is new and “Yes” if the figure is old. The subject learning memory is calculated based on both accuracy and location and z-scored across all subjects. The details of this cognitive task have previously been published (Van Erp et al., 2015).

### 2.3. Data Processing

Data obtained from fMRI were preprocessed using statistical parametric mapping (SPM12, https://www.fil.ion.ucl.ac.uk/spm/) in the MATLAB 2019 environment. The first five scans were removed for the signal equilibrium and participants’ adaptation to the scanner’s noise. We performed a rigid body motion correction using the toolbox in SPM to correct subject head motion, followed by the slice-timing correction to account for timing difference in slice acquisition. Next, the imaging data underwent spatial normalization to an echo-planar imaging (EPI) template in standard Montreal Neurological Institute (MNI) space and was resampled to 3×3×3 mm^3^. Finally, a Gaussian kernel was used to further smooth the fMRI images using a full width at half maximum (FWHM) of 6mm.

To extract reliable VSN independent components (ICs), we used the Neuromark automatic group ICA pipeline within GIFT (http://trendscenter.org/software/gift), which uses previously derived components maps as priors for spatially constrained ICA (Du et al., 2019). The Neuromark automatic group ICA pipeline was used to extract reliable, independent components (ICs) by employing previously derived component maps as priors for spatially constrained ICA. In Neuromark, replicable components were identified by matching group-level spatial maps from two large-sample HC datasets. Components were identified as meaningful regions if they exhibited peak activations in the gray matter within VSN. Although, as a previous study showed that group ICA is robust to the motion artifact (Du et al., 2016). Additional denoising and artifact removing steps prior to dynamic functional connectivity calculation were: 1) linear, quadratic, and cubic de-trending; 2) multiple regression of the 6 realignment parameters and their temporal derivatives; 3) outlier removal; and 4) low-pass filtering below a frequency of 0.15 Hz. Nine components were identified in VSN are shown in Table 1 and Step1 in Fig.1.

**Fig. 1.**
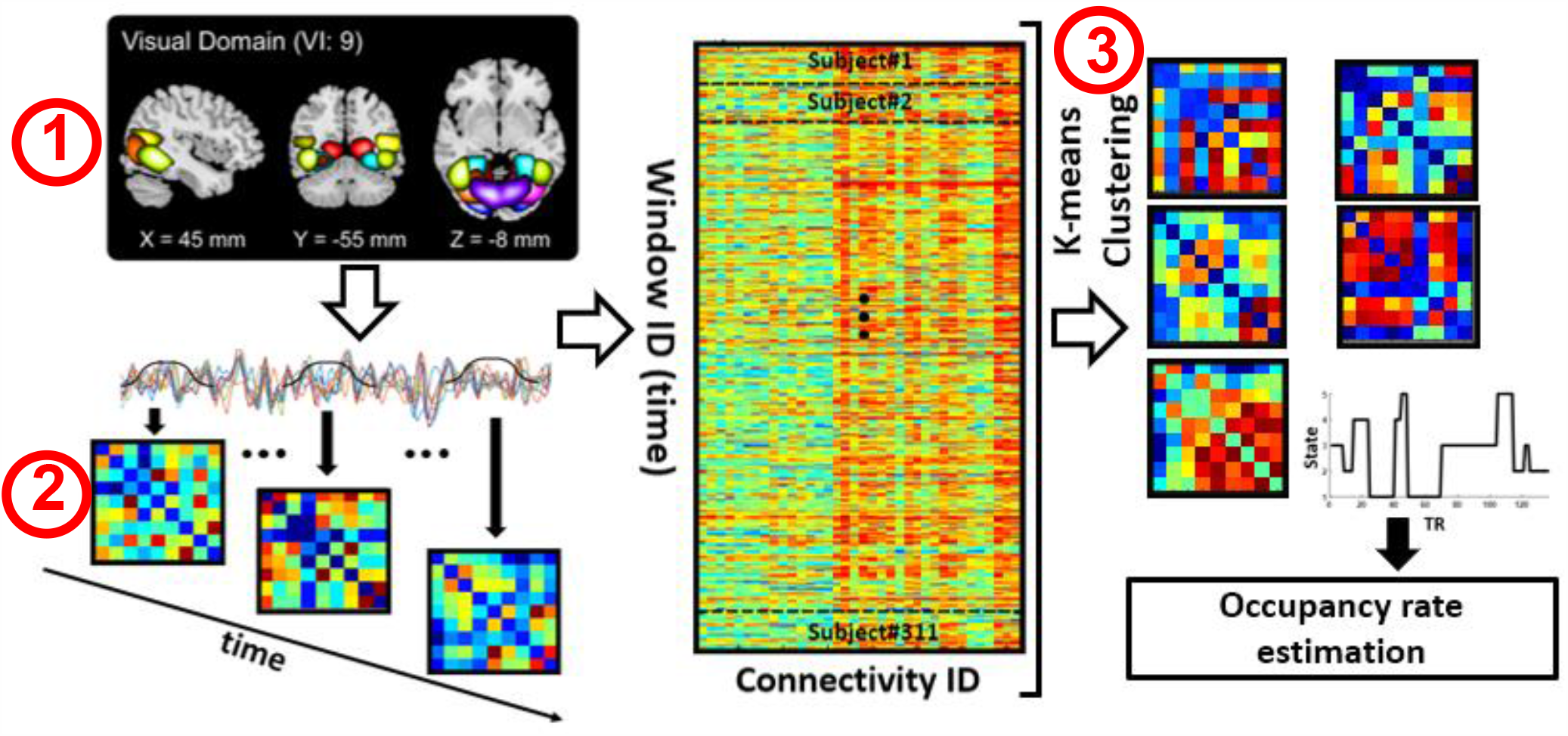
Analytic pipeline. Step1: The time-course signal of nine regions in visual sensory network (VSN) has been identified using group-ICA. Step2: After identifying nine regions in VSN, a taper sliding window was used to segment the time-course signals and then calculated the functional connectivity (FC). For each subject 137 FC matrixes were estimated and each FC matrix was a symmetric 9×9 matrix. Step3: After vectorizing the functional connectivity (FC) matrixes, we have concatenated them and then a k-means clustering with k=5 was used to group FCs to five distinct clusters for entire group and the subject-level state vector. The optimum number was k=5 based on elbow criteria. Then, occupancy rate (OCR) features, in total 5 features, were calculated from the state vector of each subject.

**Table 1.** Component Labels.

| Component name | Peak coordinate (mm) |  |  |
| --- | --- | --- | --- |
| (IC 16), Calcarine gyrus [CalG] | -12.5 | -66.5 | 8.5 |
| (IC,5), Middle occipital gyrus [MOG ] | -23.5 | -93.5 | -0.5 |
| (IC 62), Middle temporal gyrus [MTG] | 48.5 | -60.5 | 10.5 |
| (IC 15), Cuneus | 15.5 | -91.5 | 22.5 |
| (IC 12), Right middle occipital gyrus [MOG ] | 38.5 | -73.5 | 6.5 |
| (IC 93), Fusiform gyrus [FG] | 29.5 | -42.5 | -12.5 |
| (IC 20), Inferior occipital gyrus [IOG] | -36.5 | -76.5 | -4.5 |
| (IC 8), Lingual gyrus [LinG] | -8.5 | -81.5 | -4.5 |
| (IC 77), Middle temporal gyrus ([MTG] | -44.5 | -57.5 | -7.5 |

### 2.4. Dynamic functional network connectivity

To calculate dFNC, we applied a sliding window approach, as shown in Fig. 1. We used a tapered window obtained by convolving a rectangle (window size = 20 TRs = 40 s) with a Gaussian (σ = 3) to localize the dataset at each time point. Pearson correlations were calculated to measure the dFNC between 9 subnodes in VSN (figure Step1). With 9 subnodes in VSN, we estimated 36 identical connectivity features. The dFNC estimates of each window for each subject were concatenated to form a (C × C × T) array (where C=9 denotes the number of ICs, and T=137), which represented the changes in brain connectivity between ICs as a function of time (Step 2 in Fig. 1) (Calhoun et al., 2014).

### 2.5. K-means clustering and dynamic modeling

A k-means algorithm was applied to the dFNC windows to partition the data into a set of separated clusters (Allen et al., 2014; Calhoun et al., 2014; Zhi et al., 2018). The optimal number of the centroid cluster was estimated based on the elbow criterion, which is the most common method in finding the optimal value of k in k-means clustering (Damaraju et al., 2014). It is defined as the ratio of within-cluster to between cluster distance, and the objective function is minimizing this ratio. Through this procedure, we found that 5 was the optimal number by searching from k=2 to 8. The correlation was implemented as a distance metric in the clustering algorithm in 1000 iterations. The output of k-means clustering includes 5distinct states for group and state vector for each individual. A state vector shows how the network changes between any pair of states over time. In the next step, using the state vector, we found the portion of time each subject spent in each state. We called this feature the OCR of each state (Step 3 in Fig. 1). With 5 states, we had 5 OCRs for each subject.

### 2.6. Statistical analysis

We compared the functional connectivity feature of each state and OCR features between SZ subjects and HC subjects using two-sample t-tests. The connectivity feature comparison was made on 36 features, while the dFNC comparison was carried out on five OCRs features. In addition, to find the link between OCR features and the visual learning score, we used partial correlation accounting for age, gender, illness duration, antipsychotic medication doses, and scanning site. In all statistics, p values were adjusted by the Benjamini-Hochberg correction method for multiple comparisons (Benjamini & Hochberg, 1995).

## 3. Results

The following subsections highlight the main results of analyzing dFNC of VNS in SZ and HC subjects in this study. These include an overview of clinical measurement, dFNC states, the regional connectivity difference between HC and SZ in each state, and the difference between HC and SZ in OCR and its correlation with visual learning.

### 3.1. Demographic and clinical characteristics

The demographic and clinical characteristics of the participants is shown in Table 2. In SZ group, the age mean and standard deviation are 38.06 and 11.30, respectively. For HC one, the age mean and standard deviation of age were 37.04 and 10.68, respectively By using a tw0-sample t-test on all subjects (combining all sites), we did not find a significant group difference between HC and SZ in age and gender. By combining all subjects form seven sites, SZ subjects showed significant impairment in visual learning when compared to HC subjects based on the two-sample t-test (t(309)=9.22, p=1.18e-10). This difference was not significant in site 2, possibly due to their small sample sizes (see Table 2).

**Table 2.** Demographic and clinical details of subjects for each site.

|  |  | **SZ** | **HC** | **P-value** |
| --- | --- | --- | --- | --- |
| **FBIRN**<br>**Site 1** | Number | 21 | 28 | NA |
|  | Age | 30.04±8.60 | 34.78±9.40 | 0.1 |
|  | Gender(M/F) | 17/4 | 21/7 | 0.99 |
|  | Visual learning | -1.35±1.32 | 0.25±0.92 | 0.0001 |
|  | Antipsychotic medication | 304.03±144.34 (N=14) | NA | NA |
|  | Illness duration (years) | 14.45±7.35 | NA | NA |
|  | PANSS (positive) | 15.72±5.57 | NA | NA |
|  | PANSS(negative) | 14.11±3.14 | NA | NA |
| **FBIRN**<br>**Site 2** | Number | 12 | 10 | NA |
|  | Age | 44.91±11.34 | 38.10±9.39 | 0.14 |
|  | Gender(M/F) | 12/0 | 7/3 | 0.62 |
|  | Visual learning | -1.15±1.29 | -0.87±0.50 | 0.52 |
|  | Antipsychotic medication | 366.10±190.20 (N=10) | NA | NA |
|  | Illness duration (years) | 23.16±14.95 | NA | NA |
|  | PANSS (positive) | 16.90±6.70 | NA | NA |
|  | PANSS negative) | 17.20±7.39 | NA | NA |
| **FBIRN**<br>**Site 3** | Number | 24 | 27 | NA |
|  | Age | 44.41±11.90 | 42.48±12.56 | 0.57 |
|  | Gender(M/F) | 19/5 | 21/6 | 0.99 |
|  | Visual learning | -1.35±1.22 | 0.31± 1.04 | 3.49e-04 |
|  | Antipsychotic medication | 312.15±163.13 (N=21) | NA | NA |
|  | Illness duration (years) | 24.95±11.68 | NA | NA |
|  | PANSS (positive) | 16.95±4.43 | NA | NA |
|  | PANSS negative) | 16.87±5.91 | NA | NA |
| **FBIRN**<br>**Site 4** | Number | 26 | 26 | NA |
|  | Age | 36.88±12.82 | 35.23±10.56 | 0.61 |
|  | Gender(M/F) | 21/5 | 20/6 | 0.99 |
|  | Visual learning | -0.92±1.32 | -0.02±0.78 | 0.004 |
|  | Antipsychotic medication | 254.65±126.16 (N=21) | NA | NA |
|  | Illness duration (years) | 15.30±12.92 | NA | NA |
|  | PANSS (positive) | 14.29±3.77 | NA | NA |
|  | PANSS (negative) | 13.41±4.64 | NA | NA |
| **FBIRN**<br>**Site 5** | Number | 14 | 15 | NA |
|  | Age | 36.64±10.27 | 37.53±9.76 | 0.81 |
|  | Gender(M/F) | 9/5 | 10/5 | 0.99 |
|  | Visual learning | -0.95±1.21 | 0.29±1.02 | 0.0059 |
|  | Antipsychotic medication | 241.56±157.25 (N=13) | NA | NA |
|  | Illness duration (years) | 15.28±9.21 | NA | NA |
|  | PANSS (positive) | 13.14±4.46 | NA | NA |
|  | PANSS (negative) | 15±6.28 | NA | NA |
| **FBIRN**<br>**Site 6** | Number | 29 | 27 | NA |
|  | Age | 36.27±11.08 | 34.51±10.85 | 0.54 |
|  | Gender(M/F) | 17/12 | 16/9 | 0.99 |
|  | Visual learning | -0.78± 0.90 | 0.06±0.89 | 0.0009 |
|  | Antipsychotic medication | 326.18±138.77 (N=20) | NA | NA |
|  | Illness duration (years) | 15.37±12.13 | NA | NA |
|  | PANSS (positive) | 14.34±4.67 | NA | NA |
|  | PANSS (negative) | 11.93±4.06 | NA | NA |
| **FBIRN**<br>**Site 7** | Number | 25 | 27 | NA |
|  | Age | 39.56±11.80 | 37.56±10.75 | 0.52 |
|  | Gender(M/F) | 20/5 | 20/7 | 0.99 |
|  | Visual learning | -1.26±0.96 | -0.44±1.14 | 0.007 |
|  | Antipsychotic medication | 330.25±199.72 (N=20) | NA | NA |
|  | Illness duration (years) | 19.41±11.14 | NA | NA |
|  | PANSS (positive) | 16.12±5.41 | NA | NA |
|  | PANSS (negative) | 14.04±6.82 | NA | NA |
| **Total** | Number | 151 | 160 | NA |
|  | Age | 38.06±11.30 | 37.04±10.68 | 0.41 |
|  | Gender(M/F) | 115/36 | 115/45 | 0.99 |
|  | Visual learning | -1.09±1.15 | 0.001±0.94 | 1.18e<sup>-10</sup> |
|  | Antipsychotic medication | 303.26±160.01 (N=119) | NA | NA |
|  | Illness duration (years) | 18.03±12.45 | NA | NA |
|  | PANSS (positive) | 15.32±4.92 | NA | NA |
|  | PANSS (negative) | 14.32±5.42 | NA | NA |

**Note:** SZ: Schizophrenia; HC: healthy control; PANSS: Positive and Negative Syndrome Scale; M: male; F: female; NA: not applicable; all p values have been calculated using two-sample t-test.

### 3.2. Overview of dynamic FNC states

By applying k-means clustering to the dFNC across all subjects, including HC and SZ, we found five distinct clusters (states). These five reoccurring dFNC states are shown in Fig. 2. In state 1 and state 2 the middle temporal gyrus had a negative correlation with other regions of the VSN, while other regions showed a positive correlation with each other. In state 2, we observed stronger connectivity between the lingual gyrus, cuneus, and calcarine gyrus than in state 1. In state 3, the middle occipital gyrus had a strong negative connection with other regions. Compared with other states, state 4 showed more positive connectivity between inferior occipital gyrus and middle temporal gyrus. Another main characteristic of state 4 was weaker connectivity between the calcarine gyrus and other regions except for cuneus and lingual gyrus. In addition, in this state, the connectivity within the middle temporal gyrus network was stronger than that of other states. In state 5, we observed negative connectivity between the fusiform gyrus and other regions.

**Fig. 2:**
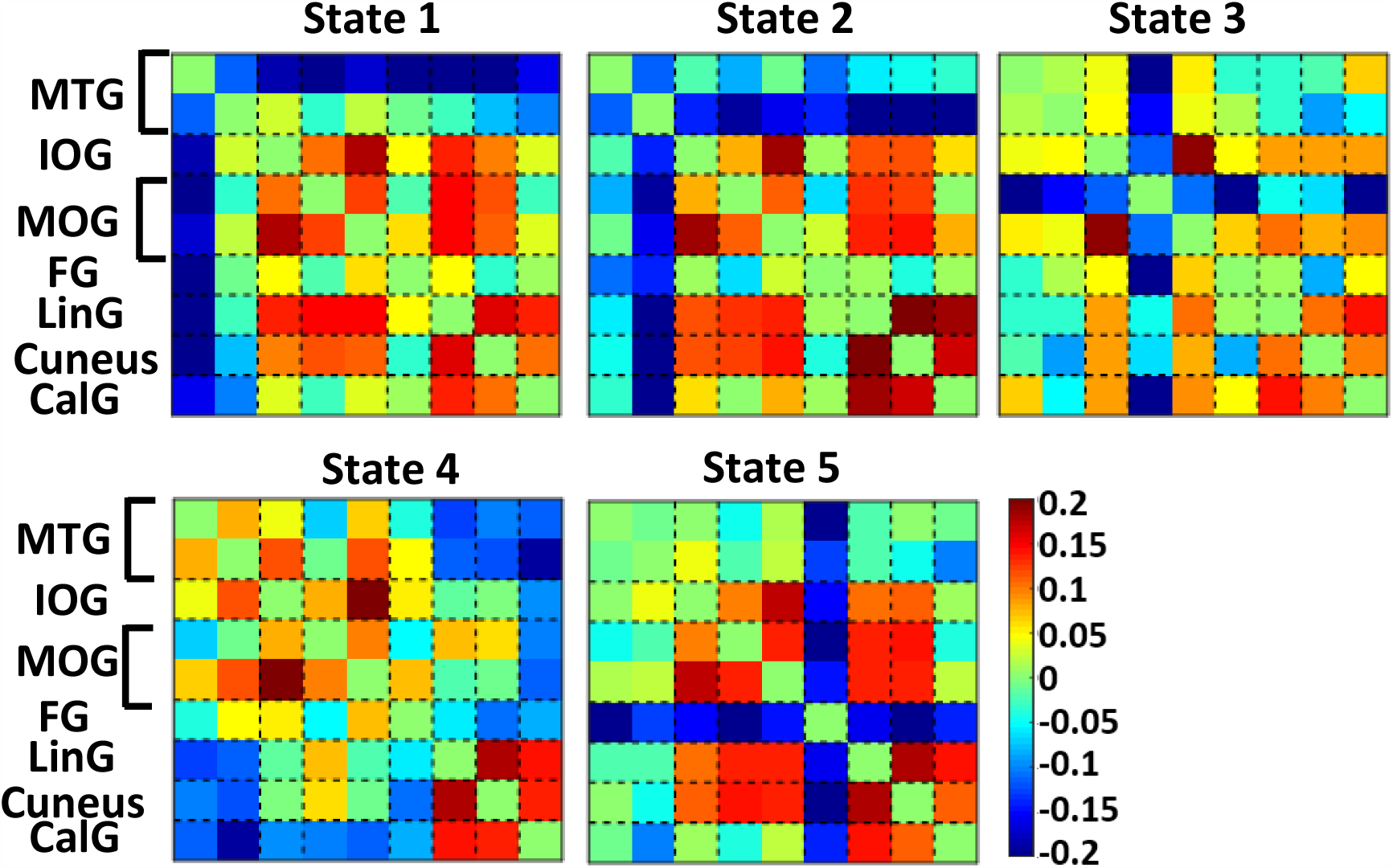
Dynamic functional network connectivity states. Five identified dFNC states using the k-means clustering method. MTG: Middle temporal gyrus, IOG: Inferior occipital gyrus, MOG: Middle occipital gyrus, FG: Fusiform gyrus, LinG: Lingual gyrus, CalG: Calcarine gyrus. State 1 and 2 showed a negative connectivity between MTG and other regions of VSN. State 4 showed the most positive correlation between MTG and other regions of VSN. In addition, this state had the maximum within-MTG conenctiivty. In state 5, FG showed a negative connectivity between FG and other regions.

### 3.3. Regional connectivity differences between SZ and HC within each state

To find differences between HC subjects and SZ subjects in each state, we used two-sample t-tests. The results are shown in Fig. 3. Significant group differences passing the multiple comparison testing are marked by asterisks (false discovery rate [FDR] corrected, q = 0.05). Interestingly, all states showed significant differences between SZ subjects and controls. In addition, we saw overlap in many of the cells showing HC vs. SZ differences. This suggests dynamics within the same sets of regions show unique fluctuations that differ significantly between HC and SZ subjects. State 3 showed the least overlap with other states. Interestingly, cuneus connectivity was significantly higher in HC subjects than SZ subjects in all states of dFNC. A similar pattern was observed in the connectivity with the middle temporal gyrus. States 2, and 3 showed fewer differences between HC and SZ in the connectivity of calcarine gyrus with other regions of VSN. The most differences between calcarine gyrus and other regions were observed in state 4. In state 5, we observed the greatest difference between HC and SZ in the connectivity of the middle temporal gyrus and other regions within the VSN. Only state 5 showed a significant difference in the connectivity of lingual and fusiform gyri between HC and SZ. It worth mentioning that we investigated the effect of illness duration and antipsychotic medication dose on the state-specific functional connectivity, and we did not observe a significant effect of illness duration and medication dose on that of the SZ subjects.

**Fig. 3:**
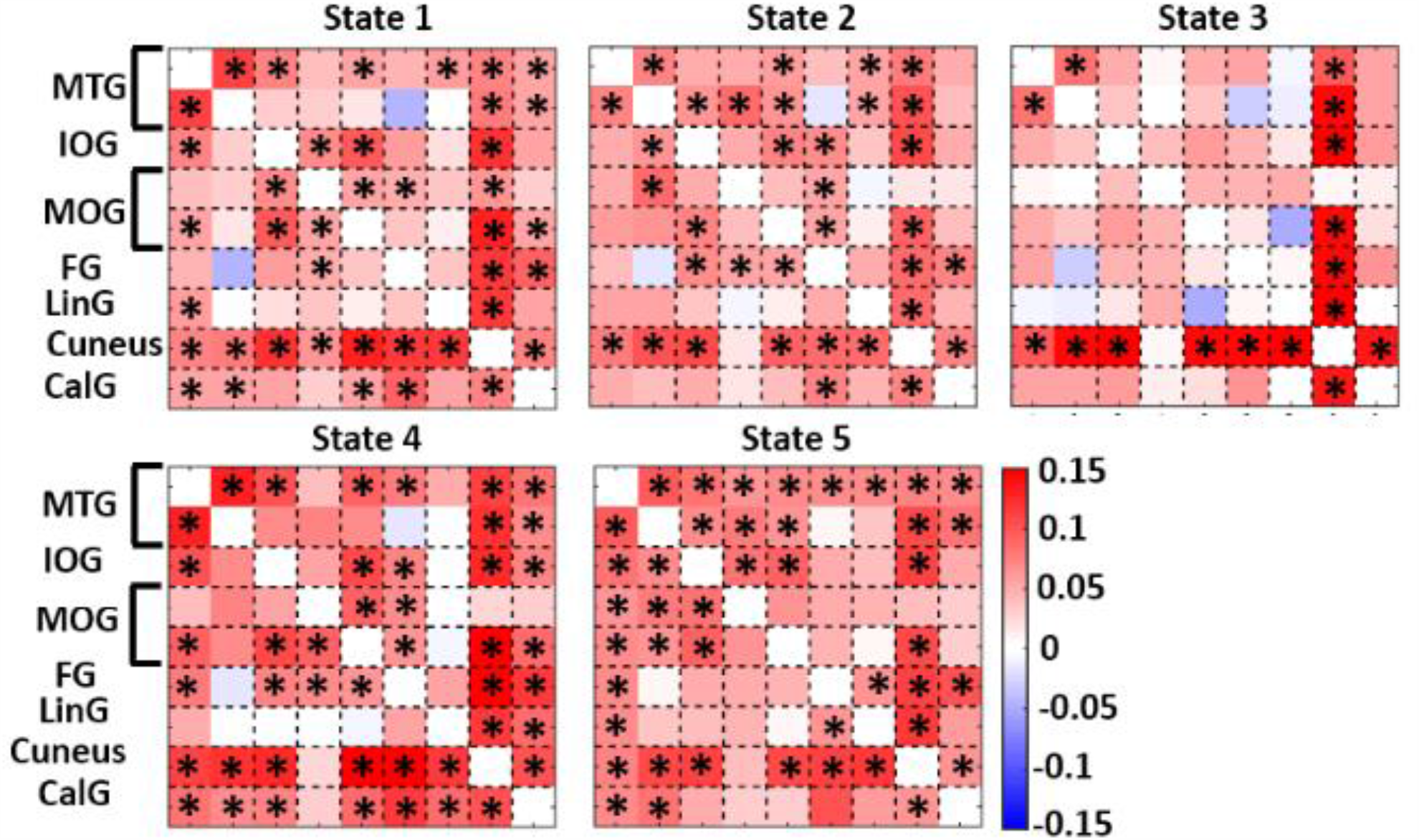
dFNC comparison between HC and SZ in each state: The visual sensory network dFNC difference between HC and SZ subjects (HC-SZ) in each state. Significant group differences passing the multiple comparison threshold are marked by asterisks (false discovery rate [FDR] corrected, q = 0.05). MTG: Middle temporal gyrus, IOG: Inferior occipital gyrus, MOG: Middle occipital gyrus, FG: Fusiform gyrus, LinG: Lingual gyrus, CalG: Calcarine gyrus.

### 3.4. Occupancy rate differences between HC and SZ and links with visual learning

Fig. 4a shows the OCR distribution of SZ and HC in each state. Using a tw0-sample t-test, we found that the OCR of state 1 was lower in HC than that in SZ (FDR corrected p=0.001). There were no significant differences in OCR between SZ and HC in any of the other states. In addition, we used a partial correlation between the visual learning score of SZ subjects and their OCR of different states while controlling for age, gender, and scanning site. The correlation between state 4 OCR showed a positive and significant correlation with visual learning memory in SZ subjects (r=0.24, corrected p=0.04. N=119), as shown in Fig. 4b.

**Fig. 4:**
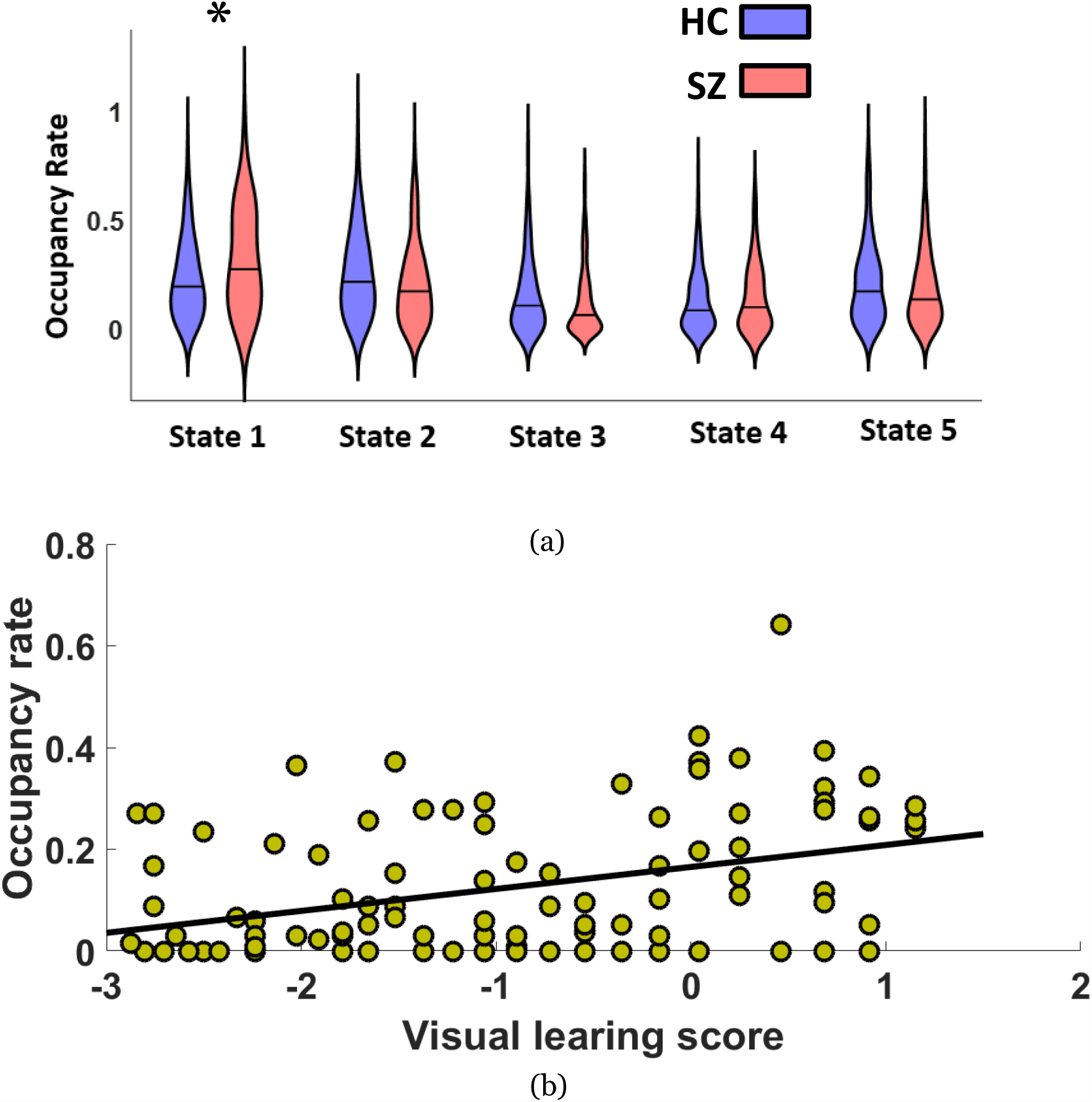
Occupancy rate difference between HC and SZ and link with visual learning: a) The difference between SZ and HC occupancy rate in different states. The occupancy rate of SZ subjects is significantly higher than that of HC subjects in state 1 (FDR corrected p=0.001). In this graph the mean value is shown by black horizontal line in each violin b) the correlation between visual learning score and OCR of state 4 in SZ subjects (r=0.24, corrected p=0.04. N=119). In this correlation, we accounted for age, gender, Antipsychotic medication, illness duration, and scanning site.

## 4. Discussion

FNC is highly dynamic, even in the absence of external input (Calhoun et al., 2014). Unlike conventional static FNC, which is obtained from the correlation within an entire time series, dFNC refers to the connectivity between any pair of brain regions within sub-portions of time series (Calhoun et al., 2014). It is likely that any cognitive deficits and clinical symptoms associated with a particular brain disorder depend not only on the strength of the connectivity between any pair of specific brain regions but also on dynamic fluctuations of connectivity among those regions. Therefore, in this study, we hypothesized that studying VSN dFNC can elucidate the mechanism of deficits in visual regions in schizophrenia.

We collected rs-fMRI scans with closed eyes in schizophrenia and healthy individuals, along with clinical, and visual learning performance data assessed by CMINDS to investigate visual network functional connectivity differences between these two groups. This is, to the best of our knowledge, the first study that explored the dynamic functional network connectivity within the VSN, including middle temporal, middle occipital, calcarine, inferior occipital, lingual, fusiform gyri, and cuneus and its association with visual learning in SZ subjects. We used a sliding window to calculate VSN dFNC over time and a k-mean clustering method to partition VSN dFNC into five connectivity patterns with unique fluctuations (states). Next, we compared the OCR for each state between SZ and HC.

By comparing FNC in different states, we found that the FNC between the middle temporal gyrus and other regions was highly negative in state 1 and state 2. However, the connectivity between this area and other regions was more positive in state 4. In addition, the connectivity among inferior occipital and middle occipital gyri was highly positive in states 1, 2, 4, and 5 with the exception of state 3. Also, middle occipital and fusiform gyrus connectivity with the rest of VSN was strongly negative in states 3 and 5, respectively. All of these observations suggest that the VSN functional connectivity is highly dynamic, even in the absence of external stimuli. This dynamic connectivity of VSN has not been explored in prior studies.

We also compared the VSN connectivity between HC subjects and SZ subjects in each state. Four states out of five states showed similar differences between SZ and HC subjects. In all states, the connectivity of cuneus in HCs was significantly higher than that in SZ subjects. This may indicate a central role of cuneus in differentiating SZ subjects from healthy individuals. It is worth noting that the cuneus, which is involved in basic visual processing, has been shown to differentiate schizophrenia from healthy subjects quite consistently. Consistent with our results, several studies reported a lower cuneus functional connectivity and altered structural properties in SZ compared to HC (Maxson & Mitchell, 2016; Yun et al., 2014; Zhou Bing, Tan Changlian, Tang Jinsong, 2010).

The correlation between the middle temporal gyrus and other regions was significantly higher in HC in all states except state 3. However, within-middle temporal gyrus correlation was significantly higher in HC subjects in all states. The middle temporal gyrus is involved in a variety of cognitive processes, including semantic memory, visual perception, and sensory integration. A study in macaques suggested the role of the middle temporal gyrus in visual connection associated with object vision. The same study showed ablation of this area causes a learning deficit in visual object discrimination and recognition (Gross, 1994). In another study, higher activation of the middle temporal gyrus was linked to perceiving facial attractiveness and expression (Vartanian et al., 2013). Several studies have reported functional and structural differences in the middle temporal gyrus between SZ and HC (Cui et al., 2018; Hu et al., 2013; Kuroki et al., 2004; Takahashi et al., 2011; Zhang et al., 2017). Middle temporal gyrus gray matter volume may be smaller in SZ subjects with first-episode schizophrenia (Kuroki et al., 2004) relative to controls. Another study reported smaller middle temporal gyrus gray matter in SZ subjects and their unaffected sibling compared to healthy subjects (Hu et al., 2013). Our findings of lower connectivity within-middle temporal gyrus regions and between middle temporal gyrus and other regions of VSN in schizophrenia, highlighted the importance of the role of middle temporal gyrus in differentiating schizophrenia subjects from healthy individuals. While the cuneus shows a consistent pattern in all states, the middle temporal gyrus contributed differently across different states. This highlights another benefit of analyzing the VSN functional network connectivity at a shorter timescale (Du et al., 2015b).

Another area showing significantly lower functional connectivity in SZ is the calcarine sulcus area. This difference was more marked in states 1 and 4. Therefore, the calcarine sulcus may play a notable role in the pathophysiology of schizophrenia, as it displayed a lower gray matter volume of calcarine gyrus in SZ subjects (Maxson & Mitchell, 2016). In addition, we found significantly lower functional connectivity of the inferior occipital gyrus and fusiform gyri in SZ subjects. Less fusiform gray matter volume and thickness have been reported in schizophrenia (Lee et al., 2002; Onitsuka et al., 2003; Van Erp et al, 2018). Also, previous literature has highlighted the role of the inferior occipital gyrus and fusiform gyrus in face processing and facial recognition (Rossion et al., 2003; Sato et al., 2012; Zhen et al., 2013). Neuroimaging studies show more activation of the fusiform gyrus in viewing face stimuli compared to nonface stimuli such as objects (Kanwishar et al., 1997). Previous studies reported abnormal face detection in schizophrenia (Chen et al., 2008). Our result can potentially highlight the role of these regions in explaining some underlying mechanisms of face processing deficits of schizophrenia and suggest further study for the role of the inferior occipital gyrus and fusiform gyrus in the face-processing deficit of schizophrenia subjects. Besides, we observed a significant difference between HC and SZ in the shorter timescale connectivity of inferior occipital, middle occipital, and fusiform gyri. It is possible that additional study of the connectivity of these regions will reveal further underlying mechanisms of visual processing deficits in schizophrenia (Rossion et al., 2003; Zhen et al., 2013).

By comparing the OCR of SZ vs. HC in each state, we found SZ subjects spent more time in state 1, showing highly negative connectivity of middle temporal gyrus with other regions of VSN. The role of dFNC temporal pattern has been reported in several disorders, including schizophrenia, major depressive disorder, and Alzheimer’s disease (Damaraju et al., 2014; Fiorenzato et al., 2019; Miller et al., 2016; Zhi et al., 2018). Our finding is consistent with previous results indicating broadly and significantly suppressed whole-brain connectivity dynamic in SZ subjects and longer periods of weak connectivity among SZ subjects relative to controls (Damaraju et al., 2014; Miller et al., 2016). A previous whole-brain dynamic connectivity analysis showed that schizophrenia subjects spend less time in a highly-connected state (Damaraju et al., 2014). Another study from our group showed an abnormal pattern in the dFNC of default mode network (DMN) by comparing state-based connectivity strength, dwell time, and between-state transition number of HC and SZ subjects (Du et al., 2015a). This study identified a SZ associated pattern in the temporal dynamics of DMN in SZ subjects by showing that they spend more time in a state with sparsely connected nodes. Also, this study demonstrated a state-specific spatial disruption within DMN by showing that the central hubs of the posterior cingulate cortex and anterior medial prefrontal cortex are significantly impaired in SZ subjects.

Finally, using the Pearson correlation between OCR and visual learning memory score in SZ subjects, we found a significant positive correlation with the OCR of state 4. Thus, SZ subjects with higher visual learning memory scores spent more time in this state. In addition, state 4 showed positive connectivity within the middle temporal gyrus and between the middle temporal gyrus with other regions, including inferior occipital, middle occipital, and fusiform gyri. This provides further evidence that the middle temporal gyrus has a significant role in dissociating SZ and HC and may also play a role in schizophrenia associated visual learning abnormalities.

Interestingly, a recent study of 467,945 subjects suggested that being born blind may be protective against schizophrenia (Morgan et al., 2018). Several studies suggested that congenitally blind people adaptively recruit and keep their visual network active for other sensory tasks, for example, high-level language or math tasks, more than healthy people (Kanjlia et al., 2016; Renier et al., 2010). In the current study, we showed more functional connectivity in the visual network of healthy people. Our findings highlight the need for further study. It would be interesting, for example, to study the dynamics of the visual system in blind individuals with this approach.

We have some limitations in this study as a previous study showed that some other factors, including dietary intake and nutritional status, drug usage, lifestyle, and even habitat, might have an effect on cognition regardless of illness pathophysiology. For example, a recent study showed that the psychopathology on a verity of measures reduced in those schizophrenia subjects who take vitamins C, E with n-3 PUFAs EPA and DHA for four months (Arvindakshan et al., 2003). Another study showed that schizophrenia subjects living in urban habitats showed more cognitive dysfunction than those who are not living in an urban environment (Arvindakshan et al., 2003). However, these factors might have an effect on healthy subjects’ cognitive function as well (Masley et al., 2017). To make sure all subjects are on the same level of intelligence, we exclude those subjects with IQ less than 75. We also believe that our large sample size (N>300) can reduce the confounds in our study.

In addition, we could not quantify whether the participant closed their eyes or stay awake during the entire time of scanning, and it is based on self-response of the subject after scanning. As a previous study showed, these might confound to the results (Agcaoglu et al., 2019). However, we assume that our dynamic functional connectivity approach would be able to separate those states in which the eyes are open, or subject are not awake from other states. That potentially proved another benefit for dynamic functional connectivity analysis, which can separate undesired states (i.e., open eyes or sleep) from the desired ones (i.e., closed eyes and awake).

Although previous studies showed that visual sensory network has a significant contribution in visual learning (Poldrack et al., 1998), other studies provide information about the contribution of other brain regions such as the hippocampus, as a part of the cognitive control network, in visual learning. In addition, a previous study showed a dysconnectivity between VSN and other brain networks in schizophrenia subjects (Damaraju et al., 2014; Sendi et al., 2020). Therefore, a prospective study on the effect of this dysconnectivity between brain networks on visual learning might provide new information about visual learning impairment in schizophrenia subjects.

In addition, VSN might contribute to other cognitive tasks, including visual working memory (Lorenc et al., 2018), speed of processing (Ho et al., 2012), visual attention (Ho et al., 2012). As a previous study showed, these cognitive tasks might be impaired in schizophrenia subjects (Javitt, 2009). Therefore, future studies are needed to explore the link between VSN dysconnectivity and the cognitive tasks mentioned above.

## 5. Conclusion

Previous studies have shown visual processing deficits in schizophrenia. Here, we investigated the dynamic functional connectivity in the resting brain’s visual sensory network. By analyzing the visual network dFNCs and clustering them into five reoccurring states, we found, as expected, that this network is highly dynamic. By comparing the VSN between HC subjects and SZ subjects in each state, we found that SZ subjects show reduced connectivity in these regions in many states. In particular, we observed that the middle temporal gyrus (in four states), cuneus (in five states), calcarine gyrus, and fusiform gyrus display decreased connection in SZ. These findings suggest a substantial dynamic disruption within the visual system in the shorter timescale in SZ subjects. In addition, we found that in transient shorter timescale estimates, SZ individuals spend more time in a state with negative functional connectivity between the middle temporal gyrus and other areas. We also found staying in a state with higher functional connectivity within and between middle temporal gyrus and other areas positively correlated with visual learning scores in SZ subjects.

Although functional connectivity can be investigated across the entire brain, the functional connectivity of fMRI visual network dynamics and their associations with schizophrenia have been understudied. With the advent of resting-state and task-associated network analyses for multiple neuropsychiatric illnesses in increasingly larger samples, identifying and comparing dynamic network connectivity may lead to novel insights into disease pathology. In sum, this work represents the first focus on functional connectivity dynamics in the visual system of schizophrenia using resting fMRI and finds substantial disruptions across multiple dynamic states. Results strongly suggest the need for a renewed focus on visual sensory network dynamics in schizophrenia both at rest and via tasks.

## Authors contribution

Mohammad S. E. Sendi developed the study, conducted data analysis, interpreted the results, and wrote the original manuscript draft. Godfrey D. Pearlson interpreted the results, edited the original draft, and provided critical review to the initial draft. Daniel H. Mathalon provided critical review to the initial draft, Judith M. Ford provided critical review to the initial draft, Theo G. M. van Erp edited the original draft and provided critical review to the initial draft. Adrian Preda edited the original draft and provided critical review to the initial draft. Vince D. Calhoun developed the study, interpreted the results, edited the original draft, and provided critical review to the initial draft. All authors approved the final manuscript.

## Acknowledgment

This work was funded by NIH grants: R01MH094524, R01MH119069, R01MH118695, and R01MH121101. Also, we thank those who helped collect this valuable data.

